# Incorporating pushing in exclusion process models of cell migration

**DOI:** 10.1101/012492

**Authors:** Christian A. Yates, Andrew Parker, Ruth E. Baker

## Abstract

The macroscale movement behaviour of a wide range of isolated migrating cells has been well characterised experimentally. Recently, attention has turned to understanding the behaviour of cells in crowded environments. In such scenarios it is possible for cells to interact mechanistically, inducing neighbouring cells to move in order to make room for their own movements or progeny. Although the behaviour of interacting cells has been modelled extensively through volume-exclusion processes, no models, thus far, have explicitly accounted for the ability of cells to actively displace each other.

In this work we consider both on and off-lattice volume-exclusion position-jump processes in which cells are explicitly allowed to induce movements in their near neighbours in order to create space for themselves (which we refer to as pushing). From these simple individual-level representations we derive continuum partial differential equations for the average occupancy of the domain. We find that, for limited amounts of pushing, the comparison between the averaged individual-level simulations and the population-level model is nearly as good as in the scenario without pushing but, that for larger and more complicated pushing events the assumptions used to derive the population-level model begin to break down. Interestingly, we find that, in the on-lattice case, the diffusion coefficient of the population-level model is increased by pushing, whereas, for the particular off-lattice model that we investigate, the diffusion coefficient is reduced. We conclude therefore, that it is important to consider carefully the appropriate individual-level model to use when representing complex cell-cell interactions such as pushing.

## I. INTRODUCTION

In humans, cell migration is an integral feature of many developmental and homeostatic mechanisms, including embryo formation [1], wound healing [2] and immune response [3]. In addition, cell migration is critical for the development and progression of pathogeneses such as cancer [4], vascular disease (e.g. atherosclerosis [5]) and chronic inflammatory diseases (e.g. arthritis [6]).

Many of the mechanisms postulated for the migration of individual cells have been well characterised in an experimental setting [7, 8]. Recently, attention has turned to studying cell migration mechanisms for cells in densely crowded environments in which cell-cell contacts are inevitable. In such environments it is possible for cells to interact mechanistically in order to facilitate movement or proliferation events. In particular, in *in vitro* experiments cells have been shown to facilitate their movement or proliferation into a region currently occupied by a neighbouring cell either crudely, by exerting direct force upon their neighbours, or, more subtly, through contact-mediated re-arrangement of a neighbouring cell’s actin-cytoskeleton leading to its dispersive migration [9].

Cells have also been shown to exert *pushing* forces on their surroundings [10]. Overcrowded groups of cells in developing epithelia have been shown to extrude cells from the epithelial sheet in order to make more room for themselves to move and proliferate into [11, 12]. Collective motion of cells, in part mediated by cell-cell pushing, has also been demonstrated to be important for normal development [13] and for the progression of pathogeneses such as cancer [9, 10]. Vroomans et al. [14] employ a cellular Potts model to infer that the pushing of de-sensitised cells by T-cells sensitive to a chemotattractant is a possible explanation for the high scanning efficiency of antigen presenting dendritic cells in the immune system.

Throughout the remainder of this paper we will refer to any contact-mediated action initiated by one cell in order to displace another to make room for itself or its progeny as a ‘push’.

Cell migration and proliferation have been modelled extensively at both the population-scale, in which deterministic partial differential equations (PDEs) are typically employed to model the density of cells [15–18], and at the cell-scale, in which each cell is modelled as an individual [19–25], often using *in silico* techniques to simulate the dynamics of the model. Both modelling regimes have their advantages and disadvantages (for a more detailed discussion of these see, for example, Baker et al. [24], Yates et al. [25]). In this paper, beginning with individual-level models (ILMs) that contain descriptions of the biological processes described above (including migration, proliferation and, for the first time, cellcell pushing), we derive population-level models (PLMs) for the evolution of the expected domain occupancy which can be thought of as being equivalent to the mean-field behaviour of the ILM in an appropriate limit.

We consider two variants of the ILM: on-lattice and off-lattice, and use a flexible master equation formalism to derive the corresponding PLM in each case. Although ILMs of cell migration and proliferation and their continuum limits have been investigated previously [19, 22], in this work we incorporate the ability of cells to displace neighbours that, in a classical exclusion process, would restrict movement or proliferation. We discover that the PDEs derived in the continuum limit from the on- and off-lattice ILMs have qualitatively different behaviour: in on-lattice models we find that pushing enhances the effective diffusion coefficient of the corresponding PDE whereas, with off-lattice models, we find that the diffusion coefficient is reduced. We provide explanations for this disparity and emphasise that it will have important ramifications for model selection when attempting to represent biological phenomena that involve cell-cell pushing.

The remainder of this paper is structured as follows. In Section II we describe, in detail, the elementary on-lattice ILM for cell-cell pushing. From this simple model we derive an equivalent population-level PDE model and, through numerical simulation demonstrate the importance of incorporating cell-cell pushing on the population-level behaviour of the cells. In Section III we incorporate more complex cell pushing mechanisms in the ILM. From these models we derive and interpret the corresponding set of PDEs that result when the appropriate continuum limit is taken. We present comparisons between the ILMs and the PLMs and comment on the causes of any disparities. We introduce the off-lattice ILM in Section IV and demonstrate the resulting PLM has some unexpected properties (in comparison to the corresponding PLM derived from the on-lattice model). We conclude in Section V with a discussion of our findings and suggestions of areas which merit further exploration.

## II. THE IMPORTANCE OF CELL PUSHING

In this section we introduce the basic on-lattice ILM and subsequently build its complexity by incorporating the ability of cells to push one another in a simple manner. Using a master equation formalism we then derive the corresponding PLM for pushing and compare the cell density generated by this model to the expected cell density averaged over several repeats of the ILM.

### A. On-lattice individual-level model

Initially we model cell migration and proliferation using a simple on-lattice, twodimensional exclusion process in which each cell is represented by a single autonomous ‘agent’. In an exclusion process, at most one agent can occupy each lattice site. We consider a square lattice (i.e. the lattice spacing is the same (Δ) in both directions) with *L*_*x*_ sites in the *x*–direction and *L*_*y*_ sites in the *y*–direction. Since an agent exclusively occupies a single lattice site, Δ can be thought of as equivalent to the diameter of the cells under consideration. The occupancy of the lattice site with index (*i*, *j*) and position (*x*,*y*) = (*i*Δ, *j*Δ) is denoted *C*(*i*, *j*). If lattice site (*i*, *j*) is occupied then *C*(*i*, *j*) = 1 otherwise *C*(*i*, *j*) = 0. We initialise *N* agents on the lattice and the occupancies of the lattice sites change in discrete time in the following manner. At each time-step, of duration *τ*, *N* agents are chosen uniformly at random, sequentially and with replacement. Selected agents attempt to move to one of their four nearest-neighbour lattice sites with probability *P*^*m*^ ∈ [0, 1] [19, 20, 26, 27]. If the site into which an agent attempts to move is occupied then that movement event is aborted. Note that sampling with replacement allows one agent to move multiple times during a single time-step and also for agents not to move at all. In what follows, we choose lattice spacing Δ = 1 and time-step *τ* = 1 noting that both time and space can be rescaled in order to deal with specific experimentally derived parameters.

In the traditional exclusion process model, if an agent attempts to move or proliferate into an occupied lattice site then that event will be aborted. In this work we relax this assumption by allowing agents to push each other out of the way in order to complete a movement or proliferation event into a currently occupied lattice site. In the most basic case (see Fig. 1 (b)) we allow an agent at position (*i*, *j*) which has chosen to move rightward into an occupied site at (*i* + 1, *j*) to push the agent at (*i* + 1, *j*) to the right into site (*i* + 2, *j*), with probability *Q*^*m*^, providing that site is unoccupied. If the site (*i* + 2, *j*) is occupied then, in this most basic case, the movement event is aborted (although we relax this condition later).

**FIG. 1.**
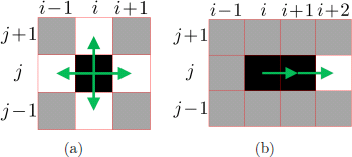
Possible movement and proliferation events in the non-pushing and basic pushing exclusion process models. Occupied sites are black and unoccupied sites are white. Sites of the lattice for which the occupancy is not important for the depicted event are shown in grey. Possible movement or proliferation directions are denoted by green arrows. (a) The selected agent at site (*i*, *j*) is free to move (or to place a daughter agent, resp.) into any of its neighbouring unoccupied sites, with probability *P*^*m*^/4 (*P*^*p*^/4, resp.). (b) The selected agent at site (*i*, *j*) has been chosen to move or proliferate to the right into an occupied site (*i* + 1, *j*). With probability *Q*^*m*^ (*Q*^*p*^ for proliferation) such that 0 ≤ *Q*^*m*^, *Q*^*p*^ ≤ 1 this agent, originally at (*i*, *j*), pushes the agent at (*i* + 1, *j*) into unoccupied site (*i* + 2, *j*) and takes its place (leaving behind a daughter agent at (*i*, *j*) in the case of proliferation).

In Fig. 2 we present snap-shot comparisons of the lattice occupancy of the exclusion process model described above, both with and without pushing and in the absence of proliferation. Some simple but informative observations can be drawn from this figure. It is evident by later times (c.f. Figs. 2 (c) and 2 (f)) that the agents that are allowed to push are more evenly spread than those that are not, with fewer large clumps of agents evident. This is to be expected as pushing agents that are clumped together are more likely to undergo successful movement events in comparison to their non-pushing counterparts, leading to the accelerated break up of such clumps. Perhaps surprisingly, the positions of the leading edge of the groups of agents are not vastly different and, after initially diverging (c.f. Figs. 2 (b) and 2 (e)), appear not to diverge more over time (*c.f.* Figs. 2 (c) and 2 (f)). This hints that, after an initial transient, the position of the leading edge is dictated primarily by diffusion events rather than by pushing. This makes intuitive sense when considering that pushing events occur more often in areas of high agent density, and are therefore less prevalent where density is low.

**FIG. 2.**
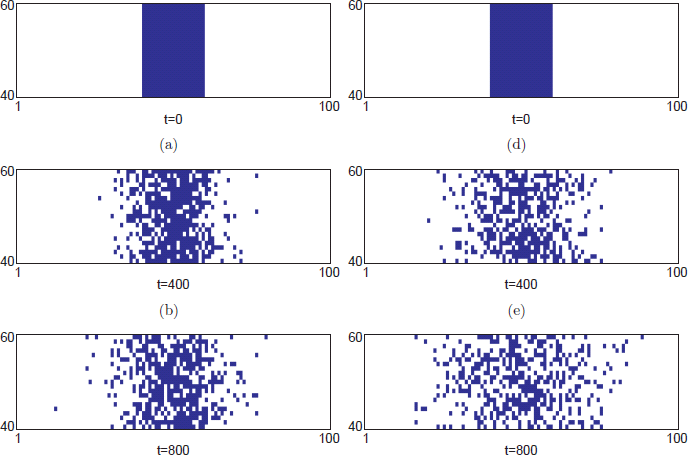
The evolution of the lattice occupancy for (a)–(c) the basic exclusion process model (*Q*^*m*^ = 0) and (d)-(f) the exclusion process model with agent-agent pushing (*Q*^*m*^ = 1). Agents are less clumped when they are allowed to push each other, although they do not appear to spread much further than in the non-pushing case. For these figures and for other on-lattice individual-level results presented later we have carried out simulations on a lattice with *L*_*x*_ = *L*_*y*_ = 100 and reflecting boundary conditions on all sides We simulate on a sufficiently large domain that any boundary effects are negligible. For clarity we only present the 21 × 100 cross-section from the middle of the domain (1 ≤ *x* ≤ 100, 40 ≤ *y* ≤ 60) at each time point. All lattice sites in the region 41 ≤ *x* ≤ 60 are initially occupied. Simulation parameters are *τ* = 1, Δ = 1, *P*^*m*^ = 0.2, *P*^*p*^ = 0.

To further quantify the difference between the spreading of the agents in the two models, we next derive the continuum equation that describes the evolution of the mean occupancy of the lattice. Comparing the effective diffusion coefficients of the model with and without pushing will provide further insight into the effect that agent-agent pushing has on the spreading of the agents.

### B. Continuum model for average occupancy

In order to derive the continuum equation for mean occupancy we first consider the probability master equation (PME) which describes the evolution of average occupancy of each site of the lattice. Let 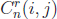 denote the occupancy of lattice site (*i*, *j*) at time *n* in the *r*^*th*^ repeat (of a total of *R* repeats) of the simulation. We define the average occupancy of site (*i*, *j*) at time *n* as

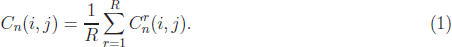

By considering the possible ways the average occupancy of site (*i*, *j*) could have changed over the course of a time-step, we can write down the following PME for the exclusion process with agent-agent pushing [28]:

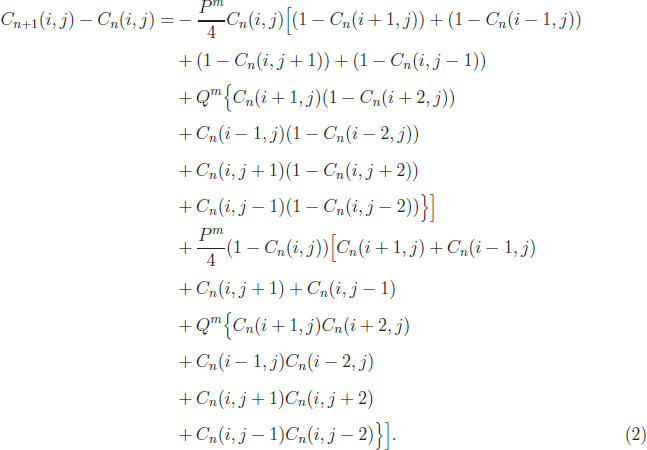

The terms that only have pre-factor *P*^*m*^ correspond to occupancy changes due to simple movement, whereas those with additional pre-factor *Q*^*m*^ correspond to occupancy changes related to movement-induced agent-agent pushing. For simplicity we ignore proliferation in this PME, but note that it is straightforward to incorporate into the PME (and the following derivation) in a similar manner.

We assume that site occupancies are independent. This simplest of moment closure assumptions can be justified in certain circumstances [19, 21, 27]. Indeed we find that for the case of basic agent-agent pushing, presented here, the PLM derived under the independence assumption agrees well with the average occupancy of the ILM (see Fig. 3 (b)). However, (as we will discover later), when more long-range agent-agent interactions are introduced the assumption begins to break down. This leads to a divergence between the mean occupancy in the ILM and the occupancy predicted by the PLM. We note that there are a variety of methods for obtaining more accurate PLMs (amongst them higher order moment closure and spatial correlation functions [29, 30]), but we do not discuss them further in this manuscript.

**FIG. 3.**
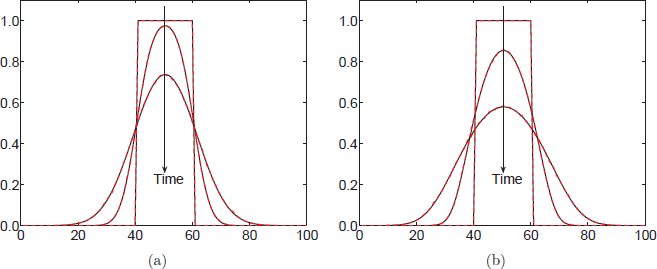
A comparison of the column-averaged density profiles of the agents in the ILM (red dashed curve) and the corresponding PDE (3) (black continuous curve) for (a) simple diffusion in the absence of pushing (*Q*^*m*^ = 0) and (b) diffusion with basic agent-agent pushing (*Q*^*m*^ = 0.5). The profiles are visualised at times *t* = 0, *t* = 50 and *t* = 200. The movement parameter in these simulations is *P*^*m*^ = 0.8. All other parameters, domain specifications, boundary conditions and initial conditions are as in Fig 2. All individual-level results are averaged over 100 repeats.

Taylor expanding the terms of the PME (2) about lattice site (*i*, *j*) and taking the (diffusive) limit of lattice size, Δ, and time-step, *τ*, both tending to zero such that Δ^2^/*τ* remains constant we obtain the corresponding PDE:

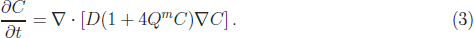

Here the diffusion constant, *D*, is given by

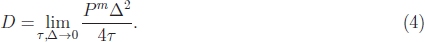

In the ILM we define the column-averaged occupancy at time *n* as follows:

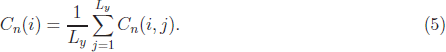

In Fig. 3 we compare the column-averaged occupancies of the ILM, both with and without pushing, to the numerical solution of the corresponding PDE (4) in one dimension [31]. The correspondence between PLM and averaged ILM is good in both cases (although marginally better for simple diffusion than for pushing (see Fig. 7 for a quantitative comparison)). As noted by considering the density profiles of the ILM the incorporation of pushing reduces peak density levels by facilitating the movement of cells away from areas of high density. This might have been predicted from equation (3) since the incorporation of agent-agent pushing increases the effective diffusion coefficient by adding a term proportional to *Q*^*m*^. This term is also density dependent, intimating that the effect of pushing will be greater when the agent density is higher and less noticeable when agent density is lower, for example at the leading edge of the profile. This density dependent phenomenon is consistent with our previous observation, from the ILM, that the positions of the leading edge of agents in the model with and without pushing are not vastly different.

In order to gain a greater insight into the possible biological effects of cell-cell pushing we now generalise the types of interactions that agents can undergo with their neighbours in the ILM and consider the effect these changes have on the resulting PDEs.

## III. EXTENSIONS TO THE PUSHING PARADIGM

It seems unreasonable, perhaps, to restrict pushing-agents to moving neighbouring agents only in their direction of movement or even for agents to be able only to move a single neighbour out of the way. We now explore the effects of relaxing these restrictions. In what follows each of the PDEs derived will be of the general form:

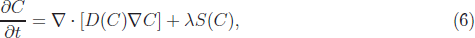

where *D*(*C*) represents an effective diffusion coefficient and *S*(*C*) a source of agents due to proliferation. Rather than writing out many different PDEs, we summarise these coefficients for each variant of the model in Table I. Note that *D* is as defined in equation (4) and *λ* is defined as follows:

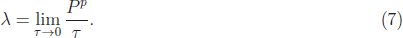

**TABLE I.** Different forms for the diffusion and source terms in the general form of the PDE (6) obtained from the different agent-agent pushing scenarios

| Pushing type |  | Coefficients |
| --- | --- | --- |
| Basic pushing | $D(C)$ | $D(1 + 4Q^m C)$ |
| | $S(C)$ | $C(1 - C)(1 + Q^p C)$ |
| Adjacent pushing I | $D(C)$ | $D\left(1 + \frac{8}{3}Q^m C\right)$ |
| | $S(C)$ | $C(1 - C)(1 + Q^p C)$ |
| Adjacent pushing II | $D(C)$ | $D\left(1 + \frac{8}{3}Q^m(C + C^2 + C^3)\right)$ |
| | $S(C)$ | $C(1 - C)(1 + Q^p(C + C^2 + C^3))$ |
| Linear pushing of multiple agents | $D(C)$ | $D\left(1 + \sum_{i=1}^K Q_i^m (i+1)^2 C^i\right)$ |
| | $S(C)$ | $C(1 - C)\left(1 + \sum_{i=1}^K Q_i^p C^i\right)$ |

### A. Pushing to adjacent positions

The first extension to the basic pushing mechanism we consider allows the agent being pushed to move into any of the free sites around it rather than simply being pushed in the direction of the movement of the pushing-agent. This can occur in two ways. In the first scenario (which we refer to as type I adjacent pushing) the pushed-agent will choose to move into one of the three potential target sites with equal probability, 1/3. If the attempted move of the pushed agent is into an occupied site then the move and the initiating push will be aborted. In the second scenario (which we refer to as type II adjacent pushing) the pushed-agent will attempt to move only into the *unoccupied* sites around it, and do so with equal probability. The agent changes how it moves based on short-range knowledge of its local environment. The three possible type II adjacent pushing movements are shown schematically in Figure 4.

**FIG. 4.**
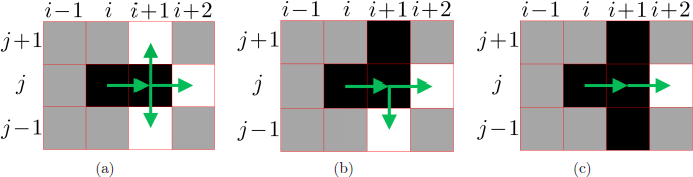
Type II adjacent pushing events in which the agent being pushed can move to any unoccupied neighbouring lattice site. The occupied sites are coloured black, whereas the unoccupied neighbouring sites are coloured white. As before, sites which do not affect the movement event are coloured grey. The selected agent at site (*i*, *j*) attempts to push to the right into the occupied site (*i* + 1, *j*). If the push is successful (with probability *Q*^*m*^) the probability with which the pushed-agent moves into an unoccupied lattice site depends upon which neighbouring sites are unoccupied. (a) The three surrounding sites about the pushed-agent are all unoccupied and the pushed-agent moves into any of them with probability 1/3. (b) Only two of the three possible sites are available and the pushed-agent moves into either of them with probability 1/2. (c) There is only one possible site for the pushed-agent to move into, which it does with certainty.

The PME for these cases become extremely lengthy and, as such, we omit them from the main text, but refer the interested reader to Section IA of the supplementary material (SM).

In Figs. 5 (a) and (b) we present a comparison of the column-averaged ILM and the corresponding PDE for type I and type II adjacent pushing, respectively. In both cases the agreement between the ILM and the corresponding PDE is good, although it is slightly better in the case of type II adjacent pushing [32].

**FIG. 5.**
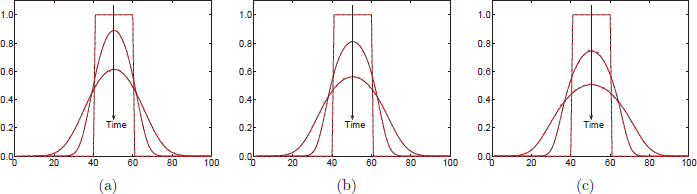
A comparison of the averaged column density profiles of the agents in the ILM and the corresponding PDE (6) with coefficients given in Table I for (a) type I adjacent pushing, (b) type II adjacent pushing and (c) multiple-agent pushing (*K* = 2). The profiles are visualised at times *t* = 0, *t* = 50 and *t* = 200. The comparison between the averaged individual results and the PDE is good in all three cases, but there is a slight under-estimation of the peak density of the ILM by the PDE for the case of multiple-agent pushing. Parameters are *P*^*m*^ = 0.8, *Q*^*m*^ = 0.5 and for (c) 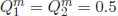. All other parameters, domain specifications, boundary conditions and initial conditions are as in Fig 2. All individual-level results are averaged over 100 repeats.

For type I adjacent pushing the peak density is not reduced as rapidly as it is for basic pushing (c.f. Fig. 3 (b) and 5 (a)). Considering the possible pushing movements from the step-function initial condition provides some insights. An agent which is one column away from the front of the initial distribution attempting to move towards the front will now only do so (by pushing an agent at the front to the right) with probability *Q*^*m*^/3, whereas in the basic pushing case it would do so with probability *Q*^*m*^. There is some compensation for the type I adjacent pushing process, in that an agent on the front which attempts to move vertically (either up or down) will now do so (displacing the neighbouring agent, whose site it moves into, out of the front in the horizontal direction) with probability *Q*^*m*^/3. In the basic pushing model these two moves would be aborted. However, we note that moves of this sort only provide a net movement of one agent in the horizontal direction in comparison to the net movement of two agents in the event of agents pushing from the column behind the front. This helps to explain why spreading is retarded in the type I adjacent pushing model.

In contrast, peak density in the type II adjacent pushing model decreases more rapidly than in the basic pushing case (c.f. Figs. 3 (b) and 5 (b)). In an analogous manner this is due to the completion of more successful pushing events; any proposed pushing event in which the pushed-agent has at least one empty neighbour will be completed with probability *Q*^*m*^ in contrast to the type I adjacent and basic pushing models in which some of these pushing events will be aborted.

### B. Pushing multiple agents in a line

The next extension we consider is to allow a pushing-agent to push up to *K* other agents in a straight line in a chosen direction (see Fig. 6). For each attempted push of *k* ≤ *K* agents we introduce a probability of acceptance 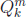.

**FIG. 6.**
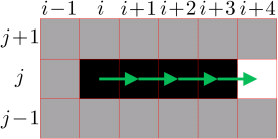
An agent attempting to move into an occupied lattice site can push up to *K* agents in a line to make room for itself. The selected agent at site (*i*, *j*) attempts to push to the right into the occupied site (*i* + 1, *j*). Sites (*i* + 2, *j*) and (*i* + 3, *j*) are also occupied. With probability 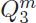 the focal agent, originally at (*i*, *j*), linearly pushes the agents blocking its path and moves into the vacated site (*i* + 1, *j*). Figure descriptions are as in Fig. 1.

The PME is, unsurprisingly, considerably more complicated than in the basic pushing case and as such we present it in Section IB of the SM. In the usual diffusive limit, upon taking the Taylor expansion about point (*i*, *j*), as before, we arrive at the PDE specified by equation (6) and the coefficients in the final row of Table I. The independence assumption that we employ in order to write down the PME (see equation (2) of the SM) becomes increasingly invalid as the number of agents that a pushing-agent can move out of the way increases. Clearly such pushing events introduce correlations between occupancies of both neighbouring and non-neighbouring lattice sites. We should not necessarily expect therefore, the comparison between the highly non-linear PDE that we derive and the averaged individual model results to be as good as in the basic pushing case. This is borne out in Fig. 5 (c) where we compare the column-averaged ILM and the corresponding PDE for linear pushing with the possibility of pushing at most two agents (i.e. *K* = 2). The agreement between the models, although slightly worse than the basic pushing case, is still at a good (see Fig. 7 for quantification). As we increase the number of agents that a single agent can push out of the way the comparison between the PLM and the averaged ILM becomes increasingly poor (See Fig. 1 of the SM).

### C. Error comparison

In order to quantify the error between the ILM and the PDE in each of the above cases, we compare averaged simulations of the ILM with the numerical solution of the PDE. Our metric of choice is the histogram distance error (HDE) [25, 33, 34]:

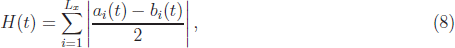

where *a*_*i*_ and *b*_*i*_ denote the values of the (normalised) column-averaged occupancies (averaged also over several repeat simulations) of the on-lattice exclusion process and numerical solution of the PDE, respectively, at lattice point *i* and time *t*.

Fig. 7 compares the evolution of the HDE for each of the above outlined cases. It is clear to see that our qualitative conclusions based on a by-eye comparison of the density profiles are borne out quantitatively by the HDE comparison. The scenario with the lowest HDE is, as expected, simple diffusion, and the scenario with the worst comparison to its ‘corresponding’ PDE is the case of linear pushing of multiple agents.

**FIG. 7.**
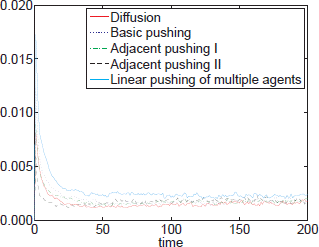
The evolution of the HDE over the time period *t* ∈ [0, 200] for each of the above presented models. Model and simulation parameters and descriptions are as in previous figures. In each case the HDE is low for the duration of the simulation. Line descriptions are as in the legend. The scenario with the best long-term correspondence to the PDE is simple diffusion (solid red line) although each of the pushing cases have similar HDEs.

We have presented results for an intermediate, representative value of the pushing parameter, *Q*^*m*^ = 0.5. However, we have also carried out comparisons for lower probabilities of successful pushing (*Q*^*m*^ = 0.1) and higher probabilities (*Q*^*m*^ = 1). The qualitative trends observed are similar for each value of *Q*^*m*^ that we considered, however, the HDE for the pushing cases was elevated when *Q*^*m*^ was increased and correspondingly reduced when *Q*^*m*^ was decreased, as might reasonably be expected (see Fig. 2 of the SM).

Similarly, although we have not presented results in which proliferation is included (for the sake of brevity and in order to clearly distinguish the effect of pushing on the diffusion coefficient) we have also carried out these simulations (results not shown) and the same qualitative results are observed for the different on-lattice pushing mechanisms.

Thus far we have considered on-lattice ILMs of cell migration. However, in reality cells do not to migrate on a regular lattice. In the next section we will relax the on-lattice assumption and explore the possibility of incorporating pushing into off-lattice models of cell migration. In particular we will explore how changing the ILM affects a corresponding PLM.

## IV. OFF-LATTICE INDIVIDUAL-LEVEL MODEL

The lattice-based models presented above provided us with a convenient formalism to incorporate agent-agent pushing into an ILM of cell migration, which was then used to derive a PLM in the form of a PDE. However, although simple to formulate, these ILMs make the important assumption that movement and proliferation events can be restricted to an artificially imposed lattice structure. This limitation is clearly an important one and it has been shown that on-lattice models can introduce artefacts which are not present in the underlying biology [35].

It makes sense, therefore, for us to consider how the effects of agent-agent pushing are altered in an off-lattice exclusion-process model of cell migration. These models are typically over-looked because of the increased complexity of their simulation and the increased mathematical complication when attempting to derive a corresponding PLM. However, recently, some excluding off-lattice models have been postulated. These models focus on considering the effects of diffusion [22, 36, 37] and proliferation [38], but none thus-far have considered the effects of agent-agent pushing.

When formulating the ILM and deriving a corresponding PLM we follow the approach of Dyson et al. [22] and Dyson and Baker [23]. We consider *N* agents of radius *R* on a line of length *L*_*x*_. Agents are initially positioned so that the gap between the centres of two adjacent agents is uniformly distributed on [2*R*, 12*R*]. In order to update the ILM we again use a random sequential update algorithm. In each time step of length *τ*, *N* agents are chosen to attempt to move. Moves are attempted with probability *P*^*m*^. A movement event consists of an agent centred at position *x* attempting to jump to *x* ± Δ. If the chosen movement would cause the moving agent to overlap with another agent, we allow a push to occur with probability *Q*^*m*^, whereby the moving agent will displace the adjacent agent enough for it to carry out movement over distance Δ. If this push is unsuccessful (with probability 1 − *Q*^*m*^) or the pushed cell would overlap another cell then the movement is aborted. See Fig. 8 for the possible ways occupancy at position *x* can change due to a single movement/pushing event.

**FIG. 8.**
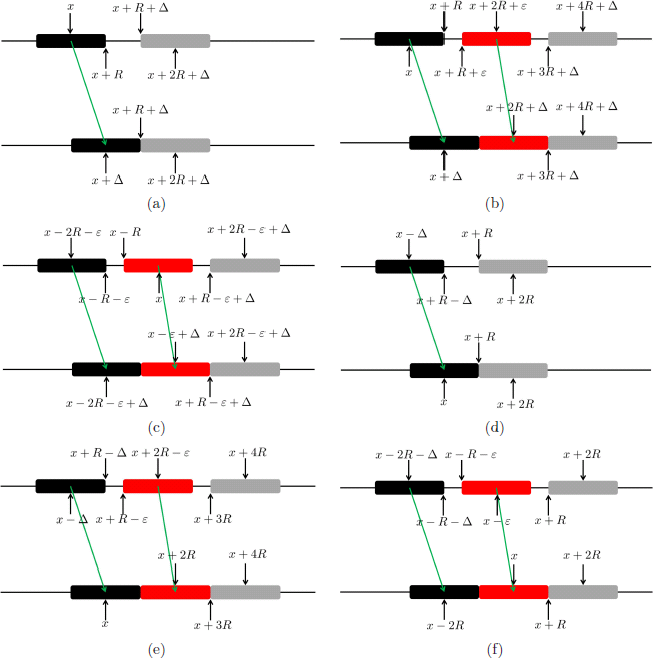
Schematics of the possible occupancy changes of position *x* due to spontaneous movement and pushing. In each panel the black agent is the agent which is moving, the red agent (if present) is the agent being pushed and the grey agent is at the closest position which another agent can be and not affect the movement or pushing of the other agents. Where it appears, 0 < ε < Δ. (a) Occupancy at *x* decreases as an agent spontaneously moves away. (b) Occupancy decreases as an agent spontaneously moves away and pushes another agent. (c) Occupancy decreases as an agent is pushed away from *x* by another agent’s movement. (d) Occupancy increases as an agent spontaneously moves to *x*. (e) Occupancy increases as an agent spontaneously moves to *x* and pushes another agent. (f) Occupancy increases as one agent is pushed to position *x* by the movement of another agent. Note that we have only shown occupancy changes due to rightward agent movement. Equivalent scenarios exist for leftward movements.

### A. Continuum model for average occupancy

We derive a PME in the same manner as for the on-lattice case, by considering the probability density functions for the positions of the agents. Let *C*_*i*_(*x*, *t*) denote the probability density function for the position of the centre of the *i*-th agent. Assuming independence of agent positions, the probability of the centre of an agent *j* (which is not agent *i*), *y*_*j*_, occupying the region [*x* + 2*R*, *x* + 2*R* + Δ) is given by

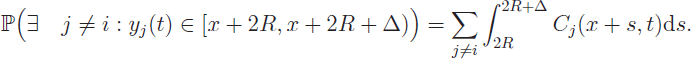

The PME can then be formulated by enumerating the possible changes in *C*_*i*_(*x*, *t*) as presented in Fig 8:

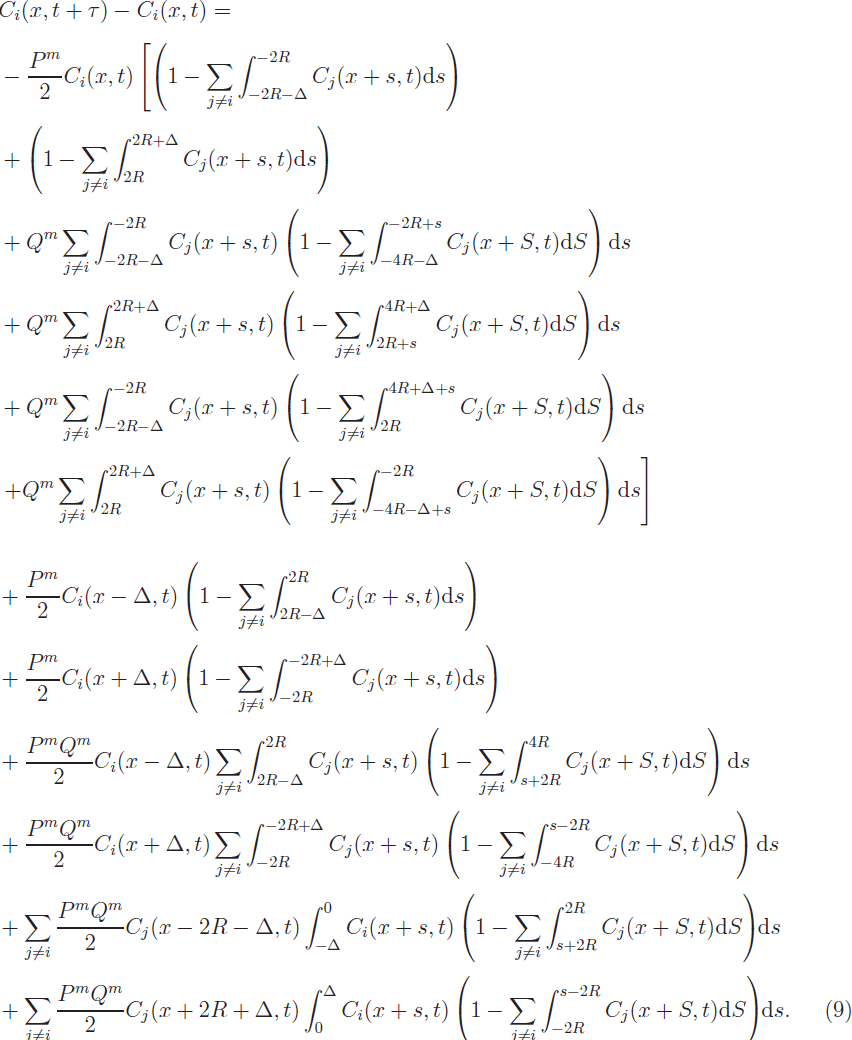

Each of the 12 lines in the PME refers to one of the panels of Fig. 8 or its leftward-moving counterpart (not shown).

We can Taylor expand these equations in *S*, provided 2*R* + Δ is small compared to the length scale on which *C* changes:

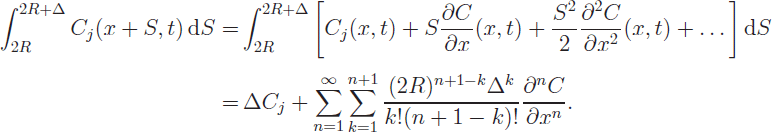

We then Taylor expand the resulting equations in *s*, in the same way, to obtain a PME free of integrals. Ignoring terms of 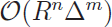 for *n* + *m* ≥ 4, leads to the following PDE for the evolution of the probability density function of agent *i* (after rearranging, dividing by *τ* and taking the usual diffusive limit):

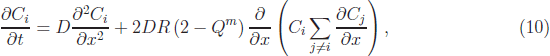

where, in analogy with equation (4),

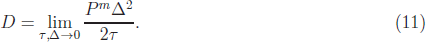

This equation for the evolution of the probability density of agent *i* is not closed, since it contains the expressions for the density of all the other agents, *C*_*j*_, *j* ≠ *i*. However, if all agent positions are initially chosen from the same distribution, then *C*_*i*_(*x*, *t*) = *C*_*j*_(*x*, *t*) ∀ *i*, *j*, so ∑_*j*≠*i*_ *∂C*_*j*_/*∂x* = (*N* − 1)*∂C*_*i*_/*∂*_*x*_. Defining the total density to be 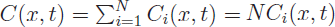 and summing equation (10) over *i*, yields:

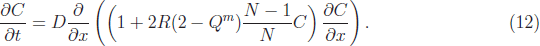

It is of comfort to note that upon setting the pushing coefficient, *Q*^*m*^, equal to zero we return to the PDE for off-lattice volume-excluding agent movement derived by Dyson et al. [22].

In order to ascertain how closely our PLM represents the average behaviour of the agents in the ILM we compare the evolution of agent density in each model. To facilitate this comparison each agent in the ILM is represented by a Gaussian kernel density function centred on its position. These averaged and smoothed individual-level simulations are compared directly to the numerical solution of equation (12) in Fig. 9. We see qualitatively that the agreement between the PDE and the averaged individual density is very good in the case of little or no pushing (panels (a) and (b) of Fig. 9, respectively). However, when pushing increases the correspondence begins to break down (see Fig. 9 (c) and (d)). In particular, the PDE over-estimates the average density of the ILM in the centre of the domain where density is high and underestimates the density when the density is lower.

**FIG. 9.**
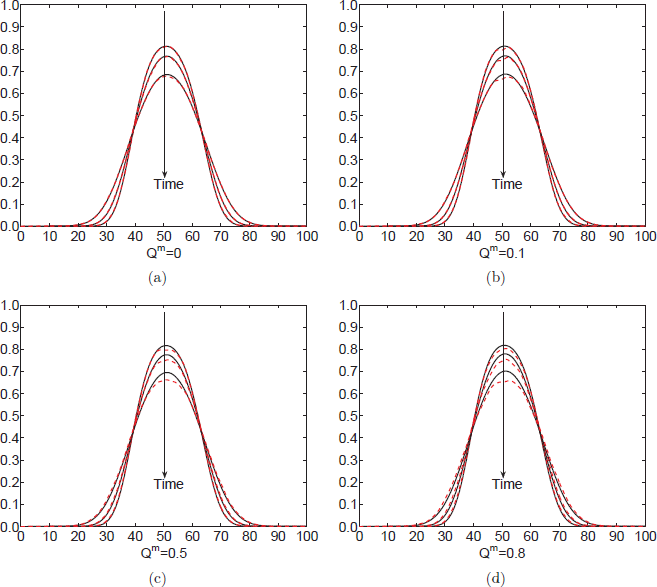
A comparison of the averaged and smoothed agent density in the one-dimensional ILM with *N* = 20 cells and the corresponding PDE (12) for a range of values of the pushing parameter *Q*^*m*^. The profiles are visualised at times *t* = 50, *t* = 100 and *t* = 200. The comparison between the PDE and the ILM reduces in quality as we increase the pushing parameter, *Q*^*m*^. Parameters for these simulations were chosen as *R* = 0.17, *d* = 0.1, *τ* = 0.04, *P*^*m*^ = 1. In each individual-level simulation agents are initialised quasi-randomly in the region [35, 65] so that no agents overlap with each other. The initial condition for the PDE is taken to be the average initial condition in the ILM. Movements which would cause an agent to leave the domain are aborted. All individual-level results are averaged over 10,000 repeats.

Previously Dyson et al. [22] and Dyson and Baker [23] have noted that agent radius, *R*, and distance moved, Δ, are key parameters resulting in changes to the diffusion coefficient. This remains true in the PDEs we derive for pushing. In particular, in Fig. 9 we do not see a distinctive change in the behaviour of the solution of the PDE as we increase the pushing parameter *Q*^*m*^. In part this can be explained by the particular choice of parameters. For our relatively small choice of *R*, volume exclusion does not change the PDE significantly from the diffusion equation. Since pushing occurs through the volume exclusion mechanism its effects on the PDE are also limited by the magnitude of *R*. A larger choice of *R* may lead to a greater ability to discern the effects of pushing in the PDE.

The evolution of the HDEs between the ILM and the PLM are shown in Figure 10. The results corroborate our qualitative conclusions from the density comparison plots. The quality of the correspondence between the two models decreases as the probability of pushing increases. We postulate that this is due, at least in part, to the break-down of our initial independence assumption with the increased propensity to push. In the off-lattice model pushing events tend to bring agents that were not previously touching into contact with each other. As such it may lead to aggregation of agents; a phenomenon which clearly breaks the independence assumption.

**FIG. 10.**
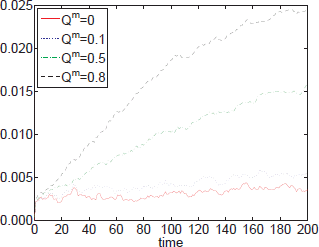
The evolution of the HDE over the time period *t* ∈[0,200] for the off-lattice model with a range of values of *Q*^*m*^. Model and simulation parameters and descriptions are as in Fig. 9. The scenarios with the best correspondence to the PDE are those with no or very little pushing (red continuous line and blue dotted line, respectively).

This ‘clumping’ phenomenon may also help to explain the particular functional form of the PDE we derive from the off-lattice ILM. The augmented diffusion coefficient has the extra density dependent term −2*RQ*^*m*^(*N* − 1)*C*/*N* in comparison to the non-pushing case. Since this term is negative the particles spreading is retarded in comparison with the non-pushing case, consistent with the idea that pushing events in the off-lattice ILM tend to gather agents together rather than disperse them. This is in stark contrast to the PDE derived from the on-lattice ILM in which pushing only serves to increase the diffusion coefficient.

## V. DISCUSSION

We have introduced a variety of on-lattice ILMs in which agent-agent pushing is explicitly incorporated and we have attempted to discern how the macroscale behaviour of models with pushing differ from their non-pushing counterparts. One quantitative way to do this is to derive the corresponding mean-field PLM and consider the solution of the PDE for different values of the pushing parameter. In each of the on-lattice cases we considered, pushing was found to augment both the diffusion coefficient and the source term (due to proliferation), in broad agreement with our findings when simulating the ILM. As we incorporated more complicated pushing mechanisms such as the linear pushing of multiple agents (rather than just one) we found that the independence assumption used to derive the PLM begins to breaks down as correlations are introduced into the ILM.

We also derived a continuum PDE from an off-lattice ILM that incorporates agent-agent pushing. Interestingly, we found that, for the off-lattice model, pushing reduced the diffusion coefficient in the corresponding PDE. In part this may be an artefact of the way we have incorporated pushing into the off-lattice model. When one cell tries to move into a region already occupied by another, it may push the obstructing cell to complete its movement event, but will remain touching the obstructing cell. The two cells that were not in contact when the movement event started are touching at the end of the movement event. As such, pushing, implemented in this manner, may lead to slower dispersal of agents corresponding to a reduced diffusion coefficient. There may be alternative off-lattice pushing mechanisms (in which pushing agents do not remain touching after a push, but rather momentum is transferred from one to the other (like billiard balls), for example) for which pushing serves to augment the diffusion coefficient, as in the on-lattice case. This serves to illustrate that it is important to accurately characterise the individual cell behaviour precisely when modelling cell migration since inaccurate characterisation can lead to qualitatively different behaviour in the resulting models.

Although we have attempted to investigate a range of different pushing mechanisms in this work, there are many questions about the modelling of cell-cell pushing which remain unaddressed. As intimated above, there are a variety of different ways to interpret cellcell pushing in both on- and off-lattice models. Some of these mechanisms may lead to increased diffusion coefficients for the PLMs corresponding to the off-lattice model or conversely decreased diffusion coefficients for the PDEs corresponding to the on-lattice model. Investigation of a variety of biologically motivated pushing mechanisms and their corresponding continuum equivalents would, therefore, be an interesting line of exploration. In addition, attempting to derive the PDE from the off-lattice ILM in higher dimensions and incorporating proliferation remain open challenges.

It is possible that cells which are pushed but have no room to move into may instigate pushes of their own in order to create space and consequently that the ‘second generation’ pushed cell instigate pushes on a third generation and so on. Although we have implemented this idea for pushing in a straight line, it may be possible to incorporate such a ‘pushing cascade’ for more complicated pushing mechanisms (such as the adjacent pushing mecha nisms) into our individual-level models. However, the increased complexity of this situation may mean that even if it is feasible to derive the corresponding continuum equation, the correlations introduced by this ‘higher-order’ pushing may render the continuum approximation a poor representation of the ILMs. In order to address this problem, and indeed the worsening correspondence between the ILM and PLM in the linear pushing model we presented in Section IIIB as *K* increases, we could consider using higher order moment closure schemes, (rather than the simple independence assumption) [39-42] or correlation functions which explicitly account for two (or more)-point distribution functions [29, 30].

Although there remains a great deal to investigate, in this work we have taken the first steps towards understanding the effects of cell-cell pushing on the macroscale migration of groups of cells. Our results have highlighted that the incorporation of pushing can be important for cell dispersal, producing qualitative changes in the corresponding macroscale PDE. However, the explicit incorporation of pushing into the ILM must be done carefully in order to capture the specific biological pushing mechanism, since different interpretations of pushing can lead to significantly different outcomes at the population level.

## ACKNOWLEDGMENTS

A.P. would like to thank the EPSRC for funding through the Oxford Systems Biology Doctoral Training Centre.

